# Myeloablation-associated deletion of ORF4 in a human coronavirus 229E infection

**DOI:** 10.1101/113423

**Authors:** Alexander L. Greninger, Gregory Pepper, Ryan C. Shean, Anne Cent, Isabel Palileo, Jane M. Kuypers, Joshua T. Schiffer, Keith R. Jerome

## Abstract

We describe metagenomic next-generation sequencing (mNGS) of a human coronavirus 229E from a patient with AML and persistent upper respiratory symptoms, who underwent hematopoietic cell transplantation (HCT). mNGS revealed a 548-nucleotide deletion, which comprised the near entirety of the ORF4 gene, and no minor allele variants were detected to suggest a mixed infection. As part of her pre-HCT conditioning regimen, the patient received myeloablative treatment with cyclophosphamide and 12 Gy total body irradiation. Iterative sequencing and RT-PCR confirmation of 4 respiratory samples over the 4-week peritransplant period revealed that the pre-conditioning strain contained an intact ORF4 gene, while the deletion strain appeared just after conditioning and persisted over a 2.5-week period. This sequence represents one of the largest genomic deletions detected in a human RNA virus and describes large-scale viral mutation associated with myeloablation for HCT.

## Introduction

Coronaviruses are positive-stranded RNA viruses with large genomes, ranging in size from 25-35kb. Although much attention has been paid to the epidemic potential and high mortality rates for MERS and SARS coronaviruses, the most common human coronaviruses (HCoV) include 229E, OC43, HKU1, and NL63 viruses ^1^. HCoV 229E and NL63 are alphacoronaviruses with genome lengths of 27-28kb, and HCoV HKU1 and OC43 are betacoronaviruses with genome lengths around 30-31kb ^2^. Few studies have been conducted on the genome evolution of the common HCoV species ^3,4^.

HCoVs enact a major burden on hematopoietic cell transplant (HCT) patients with >10% cumulative incidence of HCoV infection among HCT by day 100 of transplant ^5^. HCoV in HCT patients is associated with prolonged shedding, with a median duration of 3 weeks ^5^. Conditioning regimens for HCT usually consist of chemotherapy and irradiation that may induce genomic alterations ^6,7^. Treatment-related myeloid leukemias are well-known to be associated with radiation therapy or chemotherapeutic agents such as cyclophosphamide ^8^. Here we describe the generation of a large deletion in an RNA virus that was temporally associated with the myeloablative conditioning regimen for HCT.

## Materials and Methods

### Cell culture and qRT-PCR

The HCoV 229E type strain ATCC VR-740 was obtained and used as a positive control for cell culture. 200ul of each patient nasal swab in viral transport media (50ul 229E control/150ul 2% MEM) was inoculated onto MRC-5 cells and incubated at 37C in stationary rack for 30 minutes. After incubation, 1.2mL of 2% MEM was added to each tube and incubated at 37C. Each day tubes were read for cytopathic effect and 50uL of supernatant was stored at -80C. 50uL culture supernatants were extracted on a Roche MagNA Pure LC 2.0 and RNA was eluted in 100uL water. 10uL of RNA was used for qRT-PCR with primers targeting the HCoV polymerase gene ^9^. qRT-PCR on patient samples was performed on 200uL of nasal swab in viral transport with the same extraction and PCR conditions.

### Metagenomic next-generation sequencing and confirmatory RT-PCR with deep sequencing

mNGS was performed as described previously ^10^. Briefly, 20uL of extracted RNA was treated with DNAse I and double stranded cDNA was created using random hexamers with SuperScript III and Sequenase v2.0 (Thermo Fisher). mNGS libraries were created using one-third volume NexteraXT reactions with 16 cycles of PCR amplification. mNGS libraries from the deletion-containing specimens were prepared twice from RNA. Dual-indexed libraries were sequenced on an Illumina MiSeq over several different types of runs (1x190bp, 1x188bp, 2x94bp, 2x300bp) with consistent results. Sequencing data were Nextera adapter and quality (Q30) trimmed using cutadapt, iteratively mapped to the HCoV 229E reference genome (NC_002645) using the Geneious v9.1 read aligner, de novo assembled using SPAdes v3.9, and visualized using Geneious v9.1 and manually curated ^11–13^. The HCoV 229E reference genome and the strains sequenced in this study have an average pairwise nucleotide identity of 98%. Majority consensus genomes from mNGS were called using a minimum of 3X coverage. Reads for each specimen were also mapped to the day -12 consensus genome to look for minor variants that arose during treatment. Due to low coverage, minor variants were called with a minimum allele frequency of 10% at 11X coverage, after trimming of inversions due to Nextera artifact, variants within 10 nucleotides of end of read, and PCR duplicates.

Confirmatory gap junction RT-PCR was performed on 1uL of RNA extracted as above using 35 cycles following manufacturer’s recommended conditions (Tm 55C) using quarter-volumes of the Qiagen One-Step RT-PCR kit and primers HCoV_24034F (5´ – TTGTTGTGAATCAACTAAACTTCC – 3´) and HCoV_24808R (5´ – ACACACCAGAGTAGTACATTAAC – 3´). Deep sequencing of the day -9 and day -2 PCR amplicons was performed using 1ng of PCR product purified with the Zymo DNA Clean and Concentrator Kit-5 with 1/10^th^ volumes of the the Kapa HyperPlus kit ligation of Y-shaped adapters followed by 7 cycles of Truseq-adapter dual-indexed PCR amplification ^14^. Sequencing was performed using a 315x310bp run on an Illumina MiSeq. From the day -9 amplicon 36,880 paired-end reads were recovered, while from the day -2 amplicon 231,990 paired-end reads were recovered. Truseq-adapter and quality-trimmed using bbduk and aligned to the ORF4 intact and deleted alleles present in the day -12 and day 15 consensus genomes using the Geneious read aligner with no gaps or mismatches allowed ^15^. The methods were performed in accordance with relevant guidelines and regulations and approved by the University of Washington Institutional Review Board.

## Results

The virus analyzed was from a female in her 40s with acute myelogeneous leukemia presenting for hematopoetic cell transplant. The previous year she had undergone 4 cycles of chemotherapy (G-CLAM, G-CLA, cytarabine, and decitabine-primed MEC). Her prior month’s bone marrow evaluation showed 16% blasts by flow cytometry with normal cytogenetics. On day –12 prior to transplant, she was found to be infected by a coronavirus 229E with a cycle threshold (Ct) of 30 and persisted at the same Ct on day -9 (Figure 1A). Based on the patient’s risk of relapsing leukemia and relative rarity of HCoV pneumonitis, the decision was made to continue with HCT despite ongoing viral shedding. On days -7 and -6 she received cyclophosphamide 60mg/kg, and on days -4, -3, and -2 she received a cumulative dose of twice-daily 2 Gy total body irradiation for a cumulative dose of 12 Gy (Figure 1A). Other significant drugs the patient received during this time period included acyclovir, granisetron, micafungin, voriconazole, and sulfamethoxazole-trimethoprim. A mismatched unrelated donor peripheral blood stem-cell transplant was performed on day 0. Throughout the HCoV infection, the patient had mild nasal congestion but never required supplemental oxygen. Her course was complicated by bacteremia on day 10, thought to be due to grade 2 mucositis, and she was discharged on day 20.

**Figure 1:**
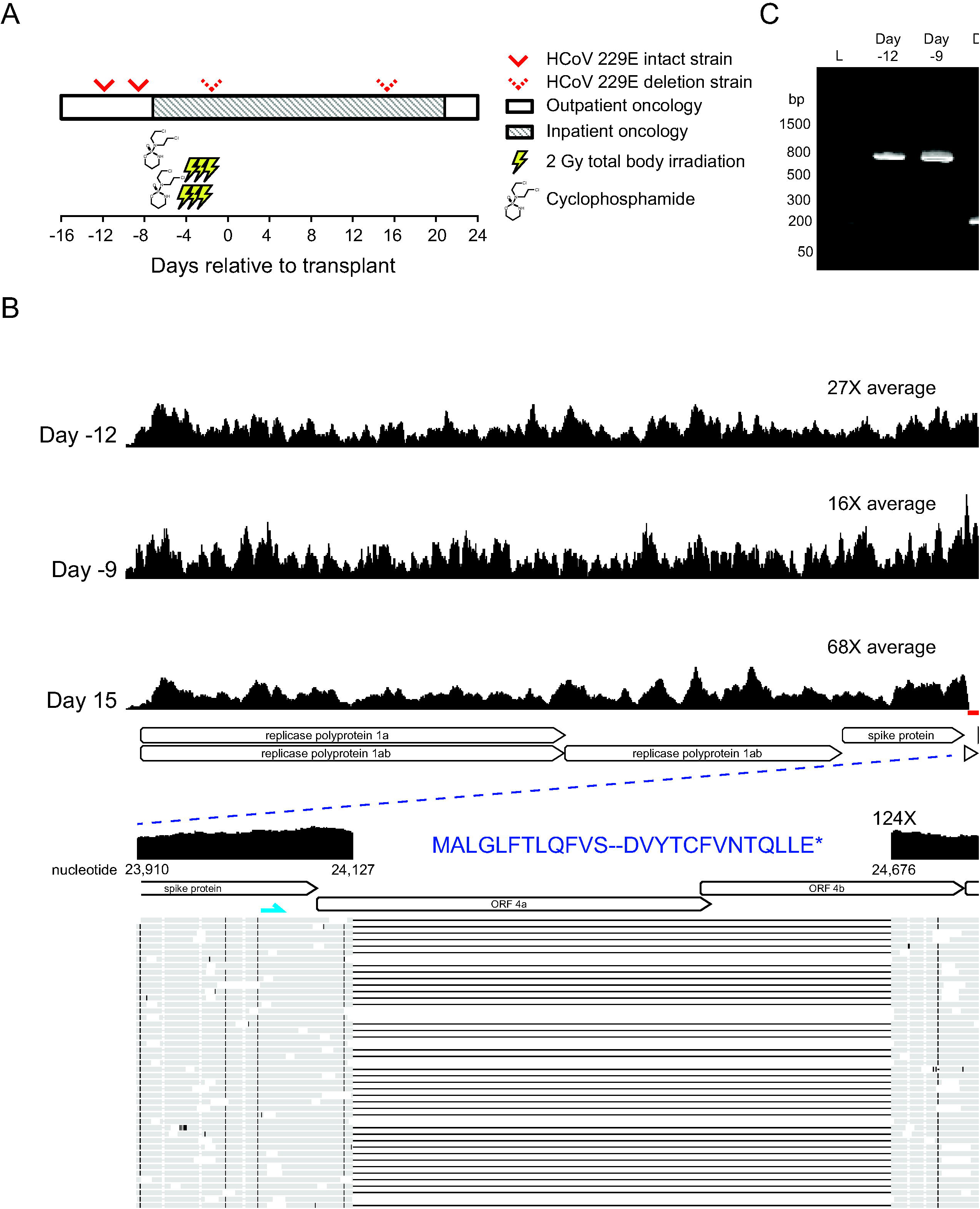
Detection of a 548-nucleotide deletion in a HCoV 229E strain associated with myeloablation. A) Depiction of the case history surrounding myeloablation and coronavirus infection. A woman in her 40s with AML was found to be infected with a HCoV 229E just before her HCT. Due to the patient’s advanced disease, the decision was made to continue with the transplant. Four successive HCoV-positive nasal swab specimens were available from the patient; two from before treatment with myeloablative cyclophosphamide and total body irradiation and two from after. All days are reported relative to HCT. B) Coverage plots of mNGS reads mapped to the reference genome for HCoV 229E (NC_002645) of three samples from the patient revealed the generation of a 548-nucleotide deletion in the ORF4a/4b gene. In addition to deleting 83% of the coding region from ORF4a/4b, the deletion resulted in a frame shift that resulted in a premature stop codon 10 amino acids before the reference genome stop codon. Primer binding sites are denoted in blue for confirmatory junction RT-PCR. C) Confirmatory junction RT-PCR of the four available HCoV-positive nasal swabs revealed the absence of the deletion before myeloablation and the presence of the deletion in specimens taken after myeloablation. NTC, no template control.

Four nasal swab samples over a four-week period were available from the patient. All qRT-PCR cycle thresholds (Ct) recovered over the four-week period were comparable, ranging between 31.3-33.5 (Table). The percent of total reads mapping to HCoV 229E ranged from 0.01-0.75%, consistent with the low concentrations of virus found by qRT-PCR (Table). mNGS of the Day 15 post-transplant specimen revealed a large deletion that encompassed 548 of the 660 nucleotides of the ORF 4 gene (Figure 1B) with average coverage of 68X across the viral genome. Repeat library preparations of the same specimen demonstrated the same deletion. The deletion was found whether mapping to the HCoV 229E reference genome or de novo assembly. No read was recovered within the area of the deletion, suggesting the presence of a viral population entirely containing the genomic deletion. mNGS of the day -12 and -9 specimens from the same patient revealed a completely intact ORF4a/4b (Figure 1B). Other than the large deletion, no consensus variants were observed between the HCoV 229E consensus genomes recovered from day - 12, day -9, and day 15 specimens. Mapping of reads from the day –12, day -9, day 15 samples to the day -12 consensus genome revealed two minor allele variants that reached >10% allele frequency and had consistent allele frequencies between the day -12 and day -9 samples. The first was a c.12332A>G, p.K4111R change in the replicase protein that went from undetectable in the day -12 and -9 specimens and was at 11.9% allele frequency in the day 15 specimen. The second was a c.675_676insU that maintained at 10.6%, 10.8%, and 8.9% allele frequency in the day -12, day -9, and day 15 specimens.

No junctional reads across the 548-nucleotide deletion were found in either the day -12 or day -9 specimen by mNGS. mNGS of the day -2 specimen also demonstrated the 548-nucleotide deletion with coverage across the genome consistent with RNA degradation (Figure S1). No consensus variants were observed in the genomic regions covered in the day -2 specimen. Confirmatory RT-PCR across the ORF 4a/4b gene demonstrated an intact gene in the pre-myeloablation specimens and a smaller PCR product consistent with the 548-nucleotide deletion in the day -2 and day 15 post-myeloablation specimens (Figure 1C). Deep sequencing of the day -9 and day -2 PCR amplicons revealed no junction reads from the day -9 specimen mapping to the day 15 deletion strain locus with 11,279X coverage at the same locus for the intact ORF4 gene in the day -12 strain (Figure S2). Conversely, only one paired-end read from the day -2 specimen mapped to the deleted portion of the ORF4 gene locus in the day -12 strain with 158,938X coverage at the ORF4 locus in the day 15 deletion strain (Figure S2).

Attempts to culture each of the patient’s specimens on MRC-5 cells proved unsuccessful. No CPE was visualized for any of the patient’s four clinical samples and HCoV 229E RNA levels in the culture supernatant were undetectable by qRT-PCR after inoculation, while viral growth was apparent in the positive control HCoV 229E ATCC type strain (Figure S3).

## Discussion

We describe metagenomic detection of a large deletion in a HCoV 229E strain temporally associated with a myeloablative regimen of cyclophosphamide and total body irradiation in an HCT patient. The ORF4-deleted virus maintained the same viral load over a 2.5 week period and was clonal by day 15 after transplant, suggesting that it was replication competent in vivo. No other majority consensus variants in the HCoV genome arose during the sampled period. One minor variant arose in the day 15 specimen that was not present in the earlier specimens, while another minor variant maintained a similar allele frequency throughout the sampled period. This rate of single nucleotide variant evolution is consistent with the molecular clock of HCoV and with a study of that found no HCoV genomic variants arose during the first month of infection ^16^. The ability of the virus to persist in vivo despite the near complete loss of ORF4 suggests that this accessory gene is completely dispensable for HCoV 229E replication and may outcompete the wild-type virus under certain conditions. Indeed, HCoV 229E type strain contains a 2-nucleotide deletion that splits ORF4 into ORF4a/4b, and the alpaca alphacoronavirus contains a 1-nucleotide insertion that also disrupts the reading frame ^17,18^. Meanwhile, all other group1b coronaviruses including all known HCoV 229E clinical strains contain an intact ORF4 ^17,19^. This is the first detection of a non-intact ORF4 in HCoV 229E associated with human infection.

The function of ORF4 in the alphacoronaviruses is unknown. The protein sequence shows no significant alignment to any protein in Protein Data Bank by HHPred ^20^. ORF4a of the HCoV 229E type strain has been suggested to be a viroporin that possessed ion channel activity ^21^. Interestingly, replacement of ORF4 in the HCoV 229E type strain with Renilla luciferase resulted in viral replication in vitro comparable to that of the wild-type strain, consistent with our in vivo data that ORF4 is unlikely to be required for replication ^22^. Taken together, these data suggest that ORF4 would make a poor antiviral target for the human coronaviruses.

Many potential mechanisms for development of the deletion exist. The chemotherapy and/or irradiation could have been directly toxic to the coronavirus genome. Cyclophosphamide is a well-known clastogen and has recently been associated with increased development of single nucleotide variations insertions and deletions ^7,23^. Alternatively, the myeloablative regime could have removed immune pressure, allowing for the loss of ORF4. It seems that in certain circumstances, deletion of ORF4 could prove a selective advantage as it was the only allele present in the day 15 specimen.

This study has several limitations. We were unable to culture any of the patient’s HCoV specimens to demonstrate ability to replicate in vitro. This is not necessarily surprising as HCoV 229E are not usually cultured in the clinical virology lab and the virus was present at low concentrations in the sample. We are also limited in accurately calling minor variants by the relatively low coverage of each viral genome. We did not attempt reverse genetics of any alphacoronaviruses to definitively show ORF4 is dispensable for in vitro replication, although the ability of the type strain to replicate despite a 2-nucleotide deletion in ORF4 and the luciferase gene replacement experiment described above are consistent with this possibility.

In summary, we describe a large deletion in HCoV 229E encompassing almost an entire gene that was temporally associated with myeloblation. Little is known about the genome evolution of HCoV 229E as only 20 genomes are currently available in NCBI Genbank. Similarly sized (500-700 nucleotide) deletions have previously been found in hepatitis D virus in patients receiving antiviral therapy ^24^. Our study is also the first demonstration of microbial evolution associated with cyclophosphamide treatment or irradiation. Further research into how chemotherapy and irradiation affect the microbiome is warranted. As mNGS is increasingly performed in the clinical lab ^25^, much intriguing biology can be unlocked from clinical specimens, providing a continuing case for the sequencing clinical microbial genomes ^26–28^.

### Data Availability Statement

These sequences are deposited in NCBI Genbank under the strain names SC677 day 15 (KY369909), SC379 day -12 (KY621348), and SC399 day -9 (KY674914). The non-human reads are deposited in the NCBI Short Read Archive under accessions SRR5809357-SRR5809372, SRR5809395, SRR5816371-SRR5816372.

### Author Contributions

ALG, GP, IP performed experiments; ALG, RCS, AC, JMK analyzed data; ALG, JTS, KRJ wrote the paper.

### Competing Interests

The authors declare they have no competing interests.

Supplementary information is available at the *Genomic Medicine* website.

**Table 1:**
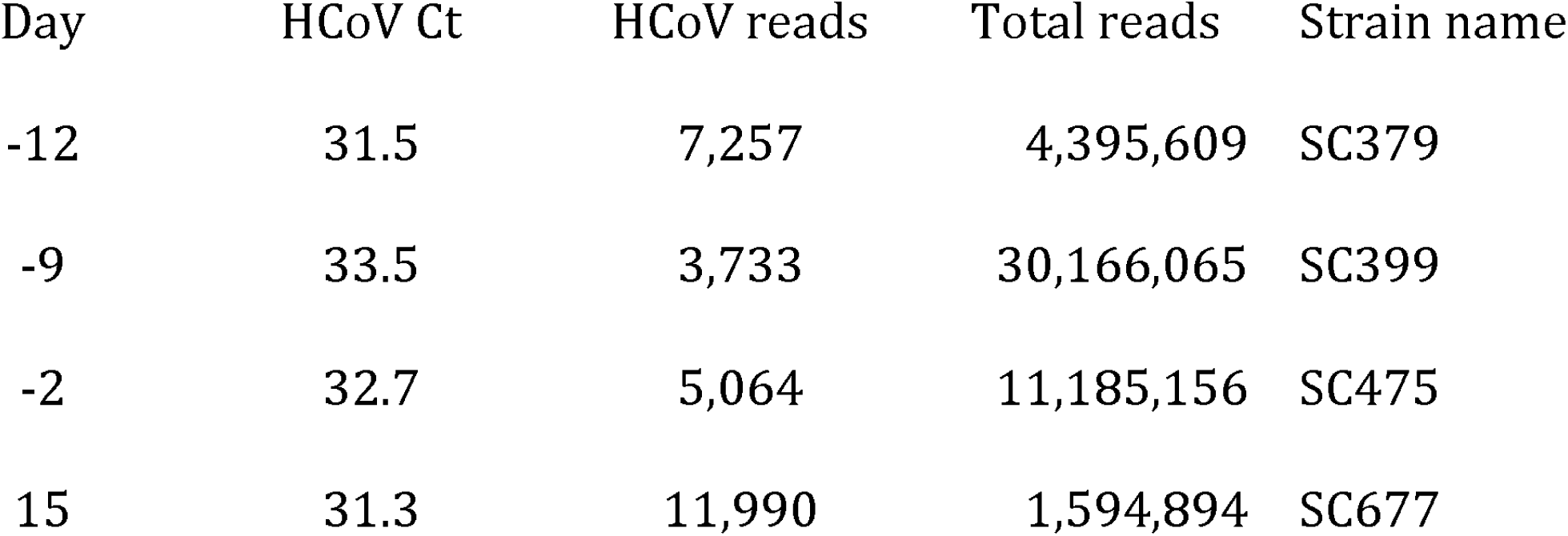
Specimens sequenced in this study. mNGS sequencing reads to HCoV 229E and total reads for each specimen are indicated, along with qRT-PCR cycle threshold.

| Day | HCoV Ct | HCoV reads | Total reads | Strain name |
| --- | --- | --- | --- | --- |
| -12 | 31.5 | 7,257 | 4,395,609 | SC379 |
| -9 | 33.5 | 3,733 | 30,166,065 | SC399 |
| -2 | 32.7 | 5,064 | 11,185,156 | SC475 |
| 15 | 31.3 | 11,990 | 1,594,894 | SC677 |

